# Precise and versatile microplate reader-based analyses of biosensor signals from arrayed microbial colonies

**DOI:** 10.1101/2022.08.12.503755

**Authors:** Fabian S. F. Hartmann, Tamara Weiß, Louise L. B. Kastberg, Christopher T. Workman, Gerd M. Seibold

**Author notes:** These authors have contributed equally to this work and share first authorship (Fabian S. F. Hartmann, Tamara Weiß). **Correspondence:** Gerd M. Seibold, Section of Synthetic Biology, Department of Biotechnology and Biomedicine, Technical University of Denmark, Søltofts Plads Building 223, DK-2800, Kongens Lyngby, Denmark.

## Abstract

Genetically encoded fluorescent biosensors have emerged as a powerful tool to support phenotypic screenings of microbes. Optical analyses of fluorescent sensor signals from colonies grown on solid media can be challenging as imaging devices need to be equipped with appropriate filters matching the properties of fluorescent biosensors. Towards versatile fluorescence analyses of different types of biosensor signals derived from arrayed colonies, we investigate here the use of monochromator equipped microplate readers as an alternative to imaging approaches. Indeed, for analyses of the LacI-controlled expression of the reporter mCherry in *Corynebacterium glutamicum*, or promoter activity using GFP as reporter in *Saccharomyces cerevisiae*, an improved sensitivity and dynamic range was observed for a microplate reader-based analyses compared to their analyses *via* imaging. The microplate reader allowed us to capture signals of ratiometric fluorescent reporter proteins (FRPs) with a high sensitivity and thereby to further improve the analysis of internal pH via the pH-sensitive FRP mCherryEA in *Escherichia coli* colonies. Applicability of this novel technique was further demonstrated by assessing redox states in *C. glutamicum* colonies using the FRP Mrx1-roGFP2. By the use of a microplate reader, oxidative redox shifts were measured in a mutant strain lacking the non-enzymatic antioxidant mycothiol (MSH), indicating its major role for maintaining a reduced redox state also in colonies on agar plates. Taken together, analyses of biosensor signals from microbial colonies using a microplate reader allows comprehensive phenotypic screenings and thus facilitates further development of new strains for metabolic engineering and systems biology.

## 1 Introduction

Phenotypic screening of microbial strain libraries for interesting genetic variants underlies many current investigations ranging from analyses of gene functions in basic research on microbial physiology to high-throughput genetic engineering of tailor-made biotechnological platform organisms ^1–4^. In the best case scenario the trait of interest is directly coupled to a phenotypic output such as growth and/or formation of a natural chromophore or fluorophore^5^ allowing the easy identification of the strains of interest ^6^. However, target phenotypes often cannot easily be detected and laborious analytical methods such as HPLC-analyses of metabolite levels would be required to identify interesting candidates. In these cases, genetically encoded biosensors have emerged as a valuable tool to facilitate high-throughput screenings. Such biosensors provide the advantage that an intracellular signal is transduced into an output signal, which can easily be measured from each strain of a library in a high-throughput manner^7,8^. Organisms naturally possess a variety of different types of sensors to monitor the intra- or extracellular accumulation of small molecules, ions, or changes in physical parameters. By applying synthetic biology tools, these properties can be harnessed to develop biosensors for high-throughput screenings ^9^. Transcription factor-based biosensors (TFBs) represent some of the most common types of biosensors applied which are often based on metabolite-sensing transcription factors. Upon interacting with effector molecules, the expression of an actuator gene, such as a fluorescent protein, is controlled ^10,11^. In contrast to TFBs, fluorescent reporter proteins (FRPs) act both as sensor and actuator. FRPs can undergo conformational changes upon interaction with a target metabolite or change in physiological state, which is subsequently accompanied by a change in their intrinsic fluorescence characteristics ^12,13^. To date, many different TFBs and FRPs are available for measuring a broad range of internal metabolites or physiological states in microbial cells ^12–18^.

Besides the selection of specific biosensors for the screening of strain libraries, the experimental set-up also represents a major determinant to be considered for an efficient screening method. High-throughput analysis of microbial libraries can be conducted in well-plates, on agar plates, via fluorescence-activated cell sorting (FACS), or droplet-based screening ^7^. Compared to FACS or droplet-based screening approaches, screenings conducted in well-plates and on agar plates significantly lowers the throughput that can be achieved. However, it provides the advantage that biosensor signals from tested strain can be directly compared under various conditions ^19^. Agar plate screens are considered as less laborious and provide a slightly higher throughput when compared to well-plate screens. However, agar plate screenings are dependent on optical readouts using camera-based imaging systems^7^. As supplier-specific filters for excitation and emission are required, the flexibility for assessing different fluorescent signals is low ^7^. Furthermore, the fixed position of light sources and camera in the imaging system affects sensitivity and might cause shadow-effects dependent of the location of the colony on the agar plate ^20^. In contrast, multimode plate readers equipped with monochromators and photomultipliers, commonly applied for well-plate screenings, offer a high flexibility, sensitivity, and easy adaptation of reader properties to the respective biosensor applied.

Despite the aforementioned challenges, we successfully applied the ratiometric FRP mCherryEA biosensor to visualize the internal pH of *E. coli* colonies growing on agar plates measured by a FUSION FX (Vilber) imaging system^8^. To adequately capture the ratiometric biosensor signals, the imaging system was equipped with two capsules (excitation laser and filter module) for excitation (530 nm and 440 nm) and a filter for capturing the emission (595 nm) of the pH-sensitive mCherry variant mCherryEA^8^.

The visualization of other biosensor signals (i.e. the redox biosensor protein Mrx1-roGFP2 with an excitation at 380 nm and 470 nm/ and an emission at 510 nm^16^) requires that the imaging system set-up is adapted to the properties of the applied fluorescent protein. Thus, applying another fluorescent protein is not readily possible if such appropriate filter modules are not available. This limitation also extends to most of the widely applied imaging systems for microbial colonies, as most cannot be equipped with varying and/or multiple fluorescence filter modules, and thus cannot be used to capture ratiometric fluorescence signals.

Recently, a standard microplate reader was applied as a tool for image-based real-time gene expression analysis using TFBs in living cells growing on the surface of solid media^21^. By scanning the surface of rectangle OmniTray plates, fluorescence signals from organisms growing on agar allowed for imaging with different resolutions (highest resolution = 360 × 240)^21^. During phenotypic screenings on agar plates, strains are typically pinned from a 96-well source plate as an array of 96, 384, or 1536 colonies on rectangular OmniTray plates ^19^. Therefore, the positions of the arrayed colonies on the agar plates are identical to the typical array of wells on conventional microplates used in microplate readers. This prompted us to test standard microplate reader systems for their applicability to assess fluorescence signals from arrayed colonies on agar plates.

In this study we demonstrate the wide applicability of microplate reader-based system to measure fluorescence signals from arrayed colonies on agar plates. This method is shown for different types of biosensors including a TFB based on LacI for regulated mCherry expression in *C. glutamicum*, promoter-based biosensors using yeast enhanced GFP (yeGFP) as fluorescent reporter in *S. cerevisiae*, and the FRP mCherryEA to assess the internal pH in *E. coli* colonies. We further show that the method developed here enables the accurate measurement of redox states for *C. glutamicum* colonies via the ratiometric sensor protein Mrx1-roGFP2 on agar plates.

## 2 Results and Discussion

### 2.1 Microplate reader-based analysis of transcription factor-based biosensors in *C. glutamicum* colonies on agar plates

TFBs are widely used in microbial physiology, metabolic engineering, and synthetic biology. Many designs for TFBs include fluorescent proteins as a reporter, which provides an easy optical readout to screen for high or low fluorescent variants from a library. Genome wide screens are often conducted via arrayed colonies on agar plates, as this approach offers a higher capacity than well-plate based screens and the comparison of different conditions^7^. However, as stated above, analyses of fluorescence signals from colonies via imaging can be challenging^22^.

To compare fluorescence imaging with microplate reader-based measurements, two model strains, *C. glutamicum* (pEKEx2_low-*mCherry*) and *C. glutamicum* (pEKEx2_high-*mCherry*), were constructed to compare fluorescence imaging with microplate reader-based measurements. The two strains were transformed using plasmids pEKEx2_low-*mCherry and* pEKEx2_high-*mCherry*, both consisting of the IPTG inducible promoter P_tac_ but differed in the strength of the ribosomal binding site (RBS) for expression of the reporter *mCherry* (Fig. 1a). To verify the different expression levels, C*. glutamicum* (pEKEx2_low-*mCherry*) and *C. glutamicum* (pEKEx2_high-*mCherry*) were cultivated in BHI liquid medium supplemented with different IPTG concentrations followed by endpoint fluorescence analysis using a microplate reader (SpectraMax). As depicted in Fig. 1b, both strains revealed an IPTG dose-dependent increase of the respective fluorescence intensity. As expected, the use of a stronger RBS in *C. glutamicum* (pEKEx2_high-*mCherry*) resulted in a higher maximal mCherry fluorescence level when compared to C*. glutamicum* (pEKEx2_low-*mCherry*). Moreover, no increased fluorescence levels were measured for the control strain *C. glutamicum* (pEKEx2) with an empty vector (Fig.1b). Analysis of fluorescence levels obtained under fully induced conditions (> 500 μM) revealed that fluorescence signals of all three strains were significantly different to each other. In presence of the highest tested IPTG concentration (5 mM IPTG), fluorescence intensities derived from the high-mCherry construct and the low-mCherry construct were approximately 50-fold and 5-fold higher when compared to the empty vector control, respectively (Fig. 1b). The results show that the constructed TFBs possess significantly different mCherry levels and both can clearly be distinguished from background fluorescence levels in liquid cultures using a microplate reader.

**Figure 1:**
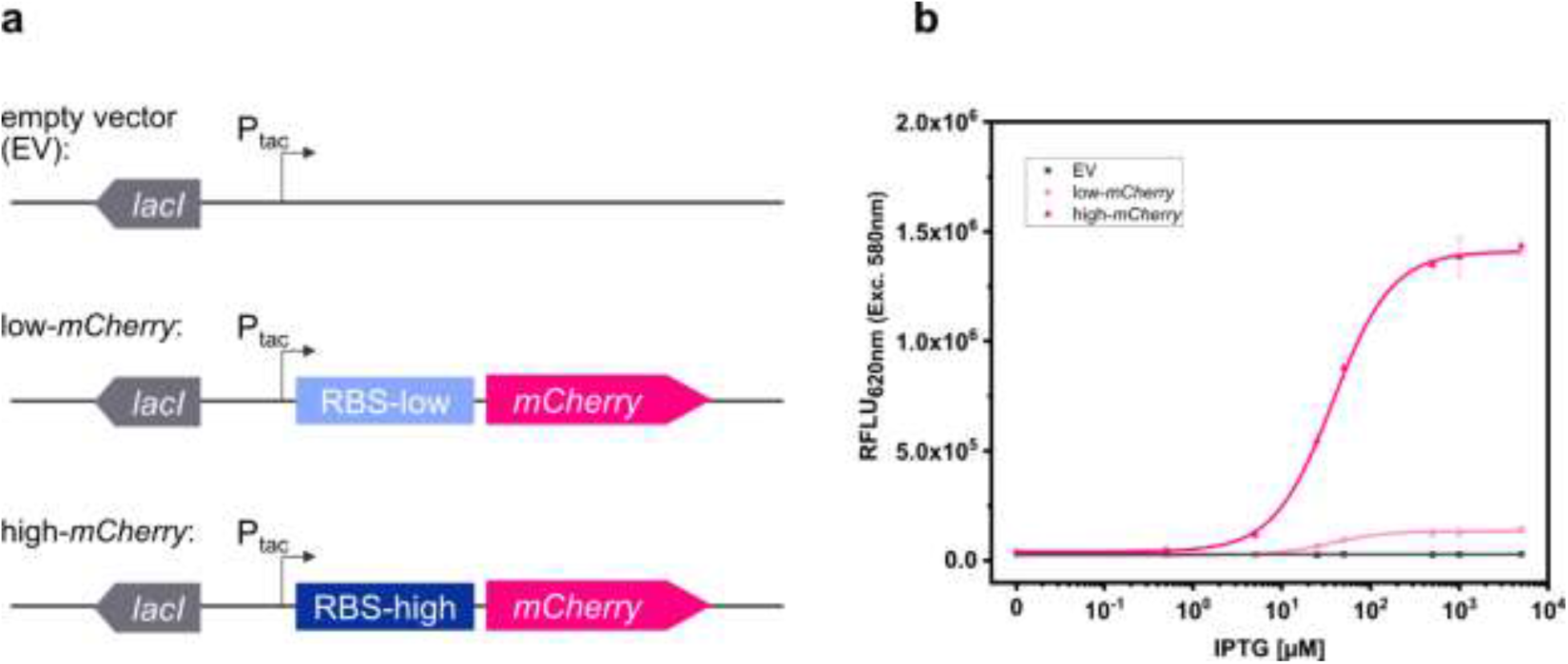
Schematic illustration of the genetic constructs for the expression platforms in pEKEx2 with different levels of mCherry translation initiation (**a**). Relative fluorescence units [RFLU] in presence of different IPTG concentrations after over-night cultivations in liquid media for *C. glutamicum* (pEKEx2) (EV: grey squares), *C. glutamicum* (pEKEx2_low_*mCherry*) (*low-mCherry:* light pink circles) and *C. glutamicum* (pEKEx2_high_*mCherry*) (high-*mCherry*: dark pink triangles) (**b**). A Hill function was fit to low-*mCherry* and high-*mCherry* data, and a linear fit was used for the empty vector (EV). Error bars represent standard deviation of four biological replicates.

To analyze the fluorescence signals derived from colonies of the three test strains *C. glutamicum* (pEKEx2), *C. glutamicum* (pEKEx2_low*_mCherry*)*, and C. glutamicum* (pEKEx2_high*_mCherry*) via imaging, the strains were spotted on rectangular OmniTray (Singer Instruments, United Kingdom) agar plates supplemented with different IPTG concentrations (0 mM to 5mM). After 48 h of incubation, the mCherry fluorescence was analyzed in a fluorescence filter equipped FUSION FX (Vilber) imaging system. In presence of 5 mM IPTG, colonies carrying the reporter pEKEx2_high_*mCherry* revealed significantly higher fluorescence intensities (mean value 8.10 × 10^2^ ± 49 RFLU) when compared to colonies carrying the empty vector control (5.11 × 10^2^ ± 43 RFLU) (Figure 2a, b). In contrast, statistical analysis via ANOVA revealed that fluorescence signals derived from colonies of *C. glutamicum* (*pEKEx2_low_mCherry*) (4.92 × 10^2^ ± 24 RFLU) are not significantly different when compared to the empty vector control (Figure 2b). To note, increased exposure times (increased from 800 ms to 1000 ms) resulted in higher fluorescence levels for both the empty vector strain, the low-mCherry as well as high-mCherry variant (Fig. S1). Albeit, fluorescence signals obtained from the high-mCherry variant reached saturation at exposure times above 800 ms, the signals detected for the low-mCherry variant were still indistinguishable from signals obtained from the empty vector control strains under fully induced conditions (Fig. S1). Significant differences in the visual detection of biosensor fluorescence are a prerequisite towards rational decision-making during screens and the selection of variants. Thus, imaging is unlikely to be suitable for all biosensors and libraries ^7^.

The accurate positioning of 96-arrayed colonies on rectangular agar plates (in a 96-well scheme) was used to measure each colony with the optics of the plate reader. To confirm the accurate positioning of arrayed colonies, we performed an absorbance scan in a 96-well scheme with 30 × 30 scans per well for an OmniTray agar plate with 96-arrayed colonies in the plate reader (Fig. S2). The absorbance scan revealed that each colony, corresponding to the respective well, was centered irrespectively of the position of the individually measured colony (Fig. S2). The exact match of the positioning allowed the plate reader to perform single point measurements similar to measurements performed using 96-well plates. Accordingly, time required for analyses of the 96-colonies can be reduced from 90 minutes (scan mode) to 20 seconds (endpoint mode). Thus, the signals obtained from the colonies represent an average value from the colony rather than capturing heterogeneity across the colony.

Based on the precision and accuracy of the colony positions, the same agar plates used for fluorescence imaging were now analyzed using a standard microplate reader. Fluorescence analysis revealed that signals detected via the plate reader for colonies equipped with the high-mCherry construct (average fluorescence of 1.98 × 10^5^ ± 5.01 × 10^3^ RFLU), as well as for colonies with the low-mCherry construct (2.13 × 10^4^ ± 7.34 × 10^2^ RFLU), were significantly different when compared to the empty vector control (4.96 × 10^3^ ± 2.88 × 10^2^ RFLU) (Fig. 2c).

In addition, we tested the dose-dependent response of the TFB carrying strains *C. glutamicum* (pEKEx2_low_mCherry and *C. glutamicum* (pEKEx2_high_mCherry) on agar plates with a series of different IPTG concentrations (0 to 5 mM IPTG; Fig. 2d). Hereby, similar dose-response curves were obtained for all strains when compared to the liquid cultures (Fig. 1b). As expected, no IPTG dependent increase of mCherry fluorescence was detected for colonies of the empty vector control strain *C. glutamicum* (pEKEx2). These results demonstrate that a microplate reader can be used for assessing fluorescence signals in arrayed colonies.

**Figure 2:**
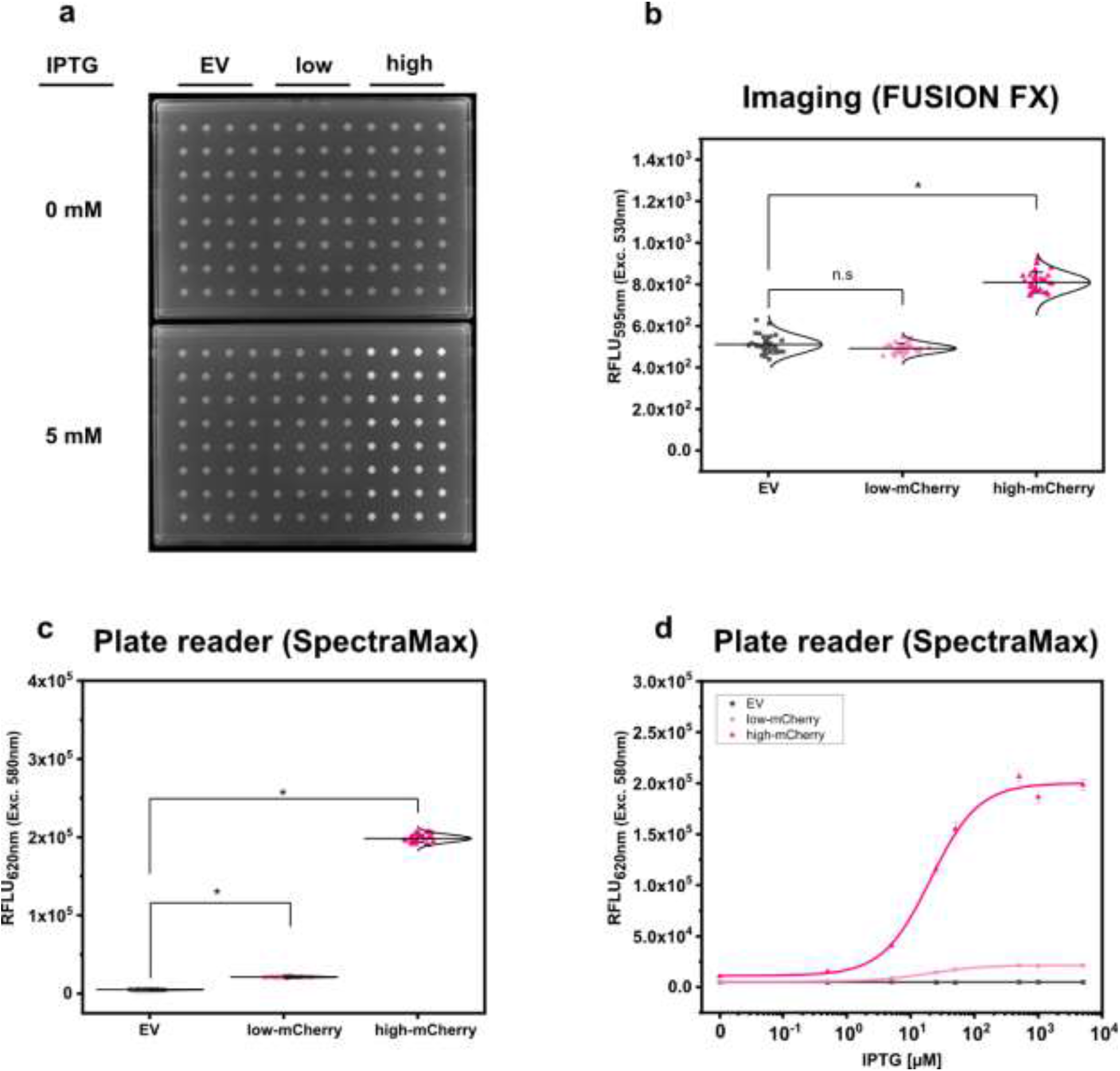
Comparison of fluorescence detection of arrayed bacterial colonies on agar plates via imaging and plate reader-based analysis. Representative fluorescence image of 32 colonies of *C. glutamicum* WT (pEKEx2) (EV: grey squares), *C. glutamicum* WT (*pEKEx2_low_mCherry*) (low: light pink circles) and *C. glutamicum* WT (pEKEx2_high_*mCherry*) (high: dark pink triangles) are spotted in absence (0 mM) or presence (5 mM) of IPTG by the use of the imaging system FUSION FX (Vilber) (**a**), the respective relative fluorescence units [RFLU] of determined for the different colonies (**b**) and the RFLU values of *C. glutamicum* colonies recorded via a plate reader (SpectraMax iD3 multi-mode plate reader (Molecular Devices LLC, U.S.A)) (**c**). RFLU values determined for arrayed colonies of *C. glutamicum* (pEKEx2) (EV: grey squares), *C. glutamicum* (pEKEx2_low_*mCherry*) (low-*mCherry*: light pink circles) and *C. glutamicum* (pEKEx2_high_*mCherry*) (high-*mCherry*: dark pink triangles) cultivated in presence of different IPTG concentrations (**d**). For sigmoidal curve fitting, Hill’s equation (low-mCherry and high-mCherry). For the empty vector (EV), a linear fit was used. For fluorescence analysis using the plate reader excitation and emission were set to 580 nm and 620 nm, respectively. Imaging of arrayed colonies on agar plates was conducted using the imaging system FUSION FX (Vilber) equipped with a capsule for excitation (530 nm) and an emission filter (595 nm). Fluorescence signals were statistically analyzed with one-way-ANOVA followed by Tukey’s test (^n.s^ p > 0.05; * p ≤ 0.001). Error bars represent standard deviation of 32 colonies.

### 2.2 Microplate reader-based system for improved fluorescence analysis of promoter-based biosensors in *Saccharomyces cerevisiae* colonies on agar plates

We next addressed the applicability of the microplate reader-based method for the analysis of fluorescence signals from the broadly used fluorescent protein GFP in colonies of the well-known model-organism S. *cerevisiae*. For this purpose, two model strains of the haploid prototrophic yeast *S. cerevisiae* CEN.PK113-7D^23^ were constructed by integration of expression cassettes with yeGFP as optical readout. Towards the development of reporter systems with different expression levels, a well-characterized weak constitutive promoter (PDA1) and strong glycolytic promoter (TDH3) were used to control varying levels of yeGFP production ^24^. Prior to analyzing the fluorescence signals from *S. cerevisiae* (background control), *S. cerevisiae* (PDA1-yeGFP) (weak promoter), and *S. cerevisiae* (TDH3p-yeGFP) (strong promoter), strains were arrayed on OmniTray agar plates.

First, fluorescence analyses of GFP signals from arrayed *S. cerevisiae* colonies was performed using the microbial colony CCD imaging workstation Phenobooth (Singer Instruments), equipped with the manufacturer’s filters for GFP fluorescence. As depicted in Fig. 3a, all colonies of *S. cerevisiae* (background) and *S. cerevisiae* (PDA1-yeGFP) (weak promoter) revealed fluorescence intensities at the same level. In contrast, significantly increased mean fluorescence intensities (1.28 ± 0.14 fold) were measured for colonies of *S. cerevisiae* (TDH3p-yeGFP) (strong promoter). However, individual signals from colonies equipped with the strong promoter still partly overlap with the values derived from the background control colonies. This means that during a random screen a high number of “GFP positive” strains would have remained undetected resulting in a low screening efficiency. Next, fluorescence signals were analyzed by the use of the imaging system FUSION FX (Vilber), equipped with a capsule and a filter for excitation at 435nm and emission at 480nm, respectively. The mean fluorescence signals derived from 96 colonies of *S. cerevisiae* (PDA1-yeGFP) (weak promoter) and *S. cerevisiae* (TDH3p-yeGFP) (strong promoter) were significantly increased by 1.30 ± 0.10 fold and 2.37 ± 0.22 fold when compared to the background control, respectively (Fig. 3b). Even though not all individual colonies of *S. cerevisiae* (PDA1-yeGFP) can explicitly be distinguished from colonies of the control *S. cerevisiae* strain, which shows background levels of fluorescence, fluorescence analysis via the FUSION FX imaging system has significantly been improved when compared to the use of an imaging workstation (Phenobooth, Singer Instruments).

As a next step, the microplate reader-based method was used to analyze yeGFP signals derived from the different promoters in *S. cerevisiae*. As described for mCherry signals in *C. glutamicum* by the use of the microplate reader SpectraMax, an average value of 6.62 × 10^4^ ± 8.59 × 10^3^ RFLU, 1.19 × 10^5^ ± 1.22 × 10^4^ RFLU and 7.46 × 10^5^ ± 8.74 × 10^4^ RFLU was determined for *S. cerevisiae* (background), *S. cerevisiae* (PDA1-yeGFP) (weak promoter) and *S. cerevisiae* (TDH3p-yeGFP) (strong promoter), respectively (Fig. 3c). This corresponds to a significant 1.80 ± 0.16 fold (weak promoter) and 11.30 ± 1.02 fold (strong promoter) increase when compared to the background control. In contrast to fluorescence imaging, all individual signals obtained for the 96 analyzed colonies harboring both the weak promoter (PDA1-yeGFP) or the strong promoter (TDH3p-yeGFP) possessed higher fluorescence levels when compared to background control colonies (Fig. 3c).

To demonstrate transferability of the microplate reader-based approach, we next analyzed fluorescence signals in arrayed colonies using a CLARIOstar^Plus^ as a different plate reader device. When compared to *S. cerevisiae*, the mean fluorescence intensity measured for all colonies equipped with the weak promoter (PDA1-yeGFP) and strong promoter (TDH3p-yeGFP) were increased 1.93 ± 0.13 fold and 12.36 ± 0.89 fold, respectively (Fig. 3d). These results are in accordance with results obtained via the SpectraMax plate reader.

Taken together, microplate reader-based analysis of colony fluorescence enables the precise and sensitive detection of fluorescence levels of reporter proteins in various microorganisms. To facilitate fluorescence analysis via this approach it is required that the colonies are arrayed in a microtiter-based format. If it is not possible to array the microbial colonies, fluorescence analysis via sensitive imaging systems equipped with adequate sets of filters, such as those used here for the FUSION FX (Vilber) imaging system, are a good alternative, although their incorporation in highly automated workflows is a challenge.

**Figure 3:**
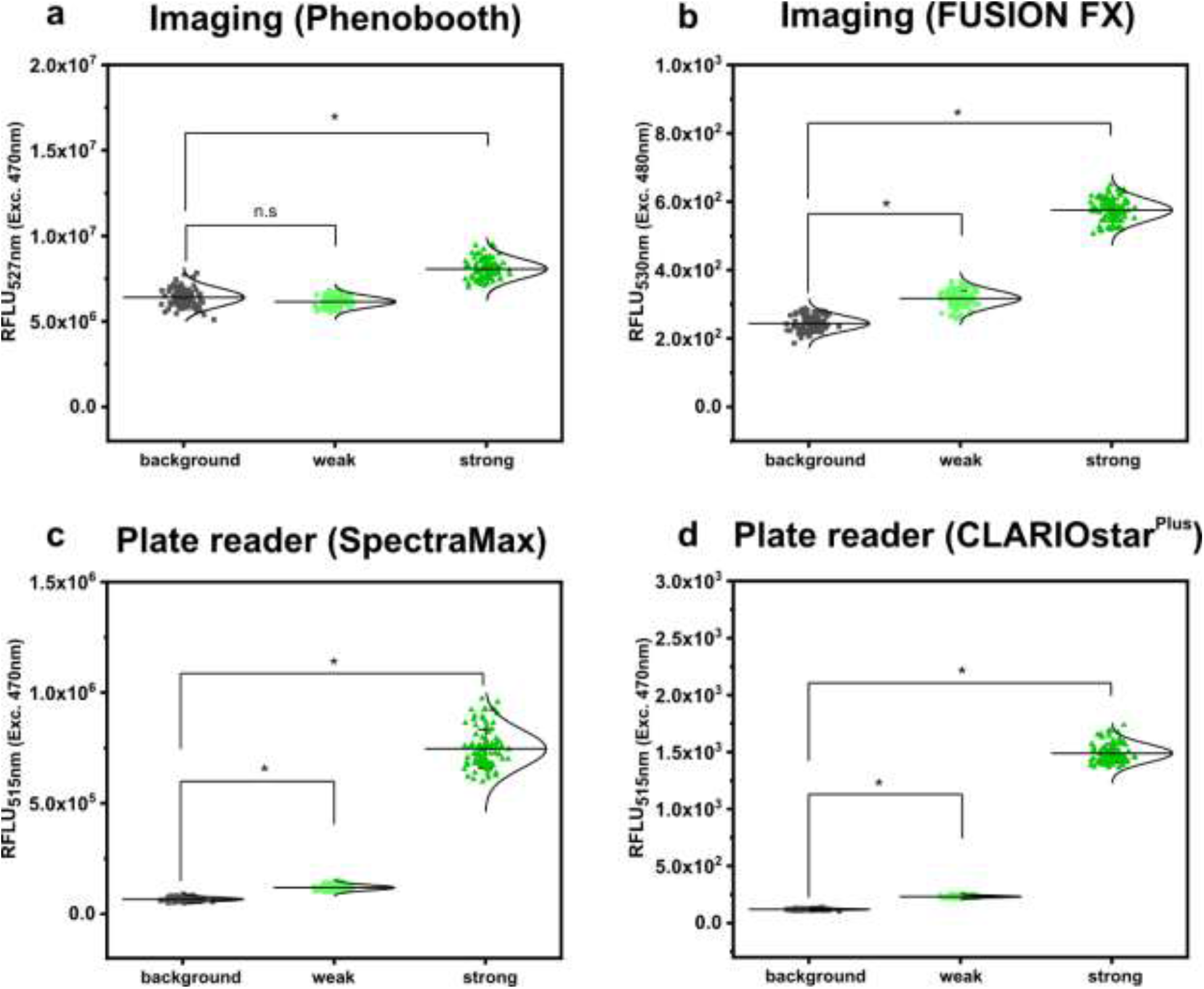
Comparison of fluorescence detection of arrayed *Saccharomyces cerevisiae* colonies on agar plates via imaging and plate reader-based analysis. Relative fluorescence units (RFLU) of 96-arrayed colonies of *Saccharomyces cerevisiae* (background; grey squares), *Saccharomyces cerevisiae* (TDH3p-yeGFP) (weak; light green circles) and *Saccharomyces cerevisiae* (PDA1p-yeGFP) (strong, green triangles) analyzed via imaging using the Phenobooth (**a**) or the FUSION FX (Vilber) (**b**). Further, fluorescence analysis was performed using the plate reader devices SpectraMax (**c**) and CLARIOstar^Plus^ (**d**). RFLU were obtained by normalizing the fluorescence intensity to the colony size (perimeter). Fluorescence signals were statistically analyzed with one-way-ANOVA followed by Tukey’s test (^n.s^ p > 0.05; * p ≤ 0.001). Mean values from 96-replicates are shown (solid horizontal line).

### 2.3 Microplate reader-based analysis of the pH-sensitive protein mCherryEA improves accuracy of internal pH measurements in *E. coli* colonies

The use of a monochromator equipped microplate reader allowed us to set optimal excitation and emission wavelength settings for the sensitive detection of mCherry and GFP signals from microbial colonies. To test if the versatility and precision of the wavelength settings via the monochromators also enables sensitive analyses of fluorescence signals from ratiometric fluorescent reporter proteins, we aimed to compare analysis of fluorescence signals from the internal pH-sensor protein mCherryEA in arrayed *E. coli* colonies via a plate reader to the recently described approach via fluorescence imaging in a FUSION FX (Vilber) imaging system^8^.

For this purpose, colonies of the strain *E. coli* MG1655 (pXMJ19_*mCherryEA*) and the empty vector control strain *E. coli* MG1655 (pXMJ19) were arrayed on SB agar plates (pH 7.0). After cultivation, 5 μL of PBS buffer supplemented with 0.05 % CTAB and with set pH values between 7.0 and 8.5 were applied onto the colonies, as recently described ^8^. At this CTAB concentration the *E. coli* cell membrane is permeabilized allowing the internal pH to become adjusted to the set external pH ^17^. The fluorescence of the colonies was analyzed using an excitation scan (400 nm – 590 nm) at an emission wavelength of 630 nm and using the positions of the 96-well measurement mode of the plate reader. As depicted in Fig. 4a, two excitation maxima at 454 nm and 580 nm were obtained for colonies of *E. coli* MG1655 (pXMJ19_*mCherryEA*), which are in accordance to the maxima observed in recently performed excitation scans of bacterial strains harboring the pH-sensitive mCherryEA protein in liquid culture^8,25^. As expected, no characteristic mCherryEA sensor signal was measured for the empty vector control *E. coli* MG1655 (pXMJ19) (Fig. 4a). When applying PBS buffer (+ CTAB) with a set pH of 7.0 to the colonies, an emission intensity of 5.8E + 06 FLU was measured at 454 nm excitation, whereas the signal obtained upon an excitation at 580 nm was higher with 6.8E + 06 FLU (Fig. 4a); when a PBS buffer (+ CTAB) with a higher pH (8.5) was applied to a colony, the fluorescence intensity obtained for an excitation at 454 nm increased (8.3E + 06 FLU) and decreased at an excitation at 580 nm (4.4E + 06 FLU). This pH-dependent response of the signals for mCherryEA at the two excitation wavelengths indicates that the ratiometric response of the biosensor protein mCherryEA in *E. coli* colonies on agar plates can be measured using the microplate reader-based system. Further, fluorescence from *E. coli* MG1655 (pXMJ19_*mCherryEA*) colonies treated with PBS buffer (+ CTAB) with set pH values ranging from 7.0 to 8.5 (Fig. 4b) were analyzed using the plate reader (Exc. 454 nm/ 580 nm) and the recently established imaging method (Exc. 440 nm/ Exc. 530 nm) ^8^. The biosensor ratios determined for the imaging method increased linear from 0.62 ± 0.02 (pH 7.0) to 1.08 ± 0.03 (pH 8.5) upon increasing the pH (Fig. 4b). A similar biosensor response was detected with the microplate reader-based method, resulting in an increase from 0.85 ± 0.03 (pH 7.0) to 1.35 ± 0.03 (pH 8.5) (Fig. 4b).

Next, 120 colonies arrayed on a rectangular agar plate were analyzed *via* the two different methods (imaging and plate reader) and the internal pH for each colony determined and visualized with a heat map, where each square corresponds to a single colony (Fig. 4c). For both methods, the mean internal pH obtained by the imaging method and the plate reader method was similar with 7.66 ± 0.13 and 7.65 ± 0.03, respectively (Fig. 4d). This agrees with the reported cytoplasmic pH of *E. coli* which is normally maintained within the range of 7.4-7.9^26–28^ and recently determined internal pH levels in *E. coli* MG1655 colonies on agar plates under similar conditions^8^. However, differences with respect to the distribution of the determined internal pH values were observed when comparing the data obtained for the two different methods (Fig. 4d). The plate reader method established here allowed us to narrow down the distribution from 7.26-7.82 (ΔpH= 0.56; imaging method) to 7.59-7.76 (ΔpH= 0.17; plate reader method), corresponding to a reduction of 0.39 pH units. As the pH is logarithmically and inversely related to the concentrations of hydrogen ions in a solution, the accuracy of the analysis is improved by 390 %. The broader distribution obtained by the imaging method might be caused by reflection artefacts from the transparent edges of the OmniTray plates. To note, the imaging method relies on a one-step excitation of the whole plate, whereas measurements performed in the microplate reader allowed us to measure colonies individually, similar to measurements performed in a 96-well microplate. Moreover, the microplate reader-based method allows operators to set excitation and emission wavelengths on demand and thus perfectly matches the two excitation maxima displayed by the sensor protein mCherryEA. In contrast, the imaging system FUSION FX (Vilber) needs to be equipped with the most suited capsules and filters available (Exc. 440 nm & Exc. 530 nm), which often do not match the properties of the fluorescent protein^25,29,30^.

**Figure 4:**
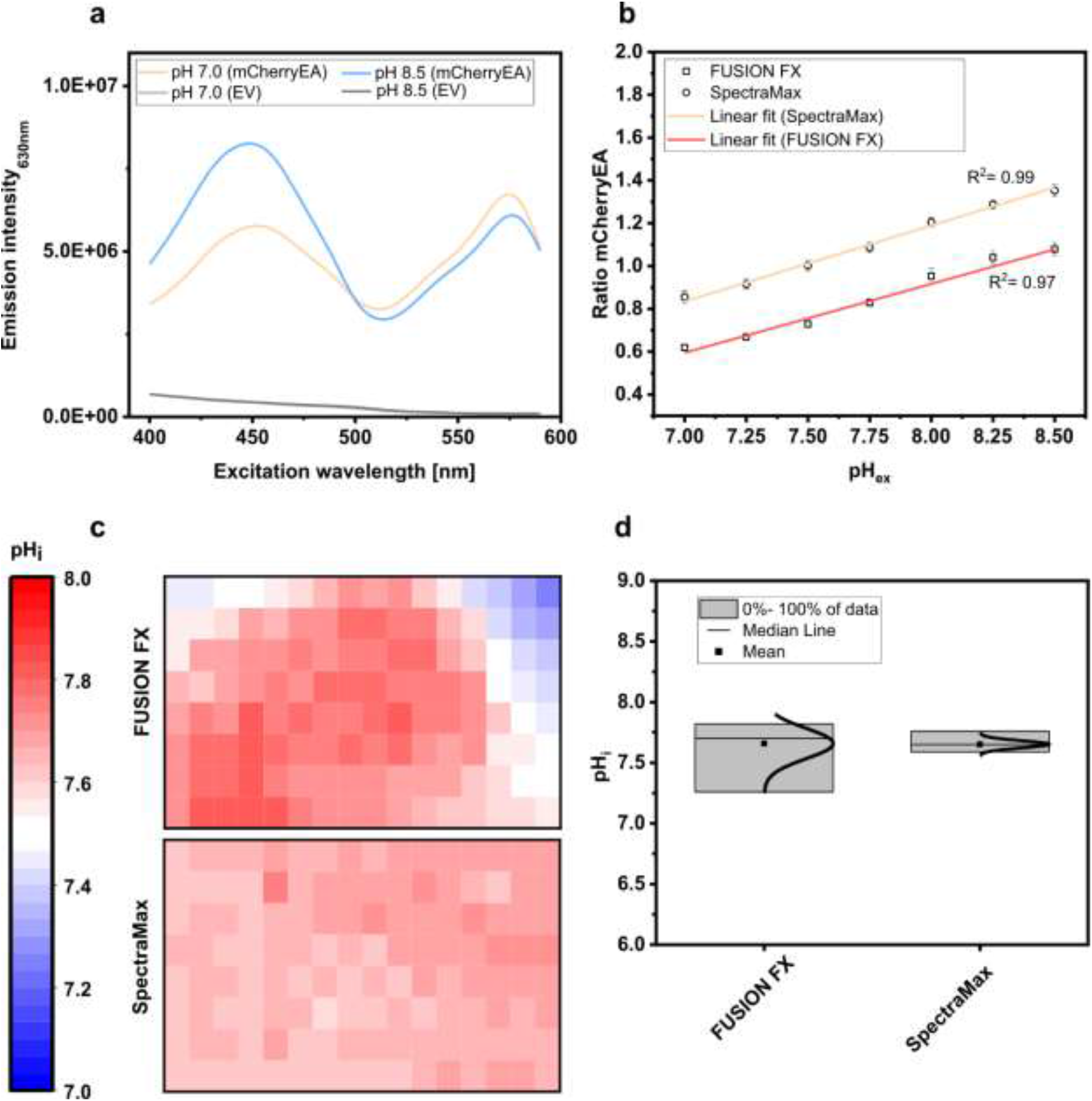
Excitation scan (400 nm – 590 nm) recording the emission intensity at 630 nm in colonies of *E. coli* MG1655 (pXMJ19_*mCherryEA*) (mCherryEA) and the empty vector control strain *E. coli* MG1655 (pXMJ19) (EV) after applying 5 μL PBS buffer supplemented with cetyltrimethylammonium bromide (CTAB; final concentration 0.05 % (w/v)) with a set pH of 7.0 and 8.5 (**a**) and ratiometric biosensor signals obtained by dividing the emission intensity at 454 nm by 580 nm upon applying PBS buffer with pH values ranging from 7.0 – 8.5 (**b**). Calculation of the internal pH for biosensor signals recorded from 120 colonies either *via* imaging using the FUSION FX imaging system equipped with two capsules (Exc. 440 nm & Exc. 530 nm) and one emission filter (Em. 595 nm) as recently described in Hartmann et al. 2022^8^ or with a microplate reader system (SpectraMax) (**c**) and data distribution with median and mean values obtained by the two different approaches (**d**). Error bars represent standard deviation from at least three replicates.

### 2.4 Mycothiol is required to maintain a reduced environment in *C. glutamicum* colonies growing on agar plates

The improved accuracy using the plate reader-based biosensor analysis represents a significant step forward with respect to screening approaches requiring high sensitivities. Moreover, the high flexibility brought by the monochromatic technology of the plate reader device easily adapts to other fluorescent reporter proteins without causing additional costs and effort to equip imaging devices with appropriate filters. Thus, we next tested this novel technique by applying a ratiometric redox biosensor protein called Mrx1-roGFP2 in *C. glutamicum* colonies.

The abundant low molecular weight (LMW) mycothiol (MSH) functions to maintain the reduced state of the cytoplasm and represents the main non-enzymatic antioxidant in high-GC Gram-positive bacteria, such as the industrial amino acid producer *C. glutamicum* ^31–33^. Recently, the genetically encoded redox biosensor protein Mrx1-roGFP2 was successfully applied in *C. glutamicum* WT and the MSH-deficient mutant strain *C. glutamicum ΔmshC* to monitor dynamic redox changes in liquid cultures^16,34^. Mutant strains lacking MSH have revealed high susceptibility towards oxidative stress resulting in an impaired growth behavior when cells were exposed to artificial oxidants in shaker-flasks^32^. In absence of artificial oxidants, growth of the MSH-deficient mutant strain proceeded similar to the wild type strain^16,32,34^, even though biosensor measurements in the mutant strain revealed an oxidative redox shift ^16,34^. Detecting intracellular changes (i.e. oxidative stress) prior to the occurrence of a growth defect is an important advance for the development of highly sensitive sensor-based screening approaches. This prompted us to test the analysis of intracellular redox states in arrayed colonies of *C. glutamicum* WT and a mutant strain lacking MSH using the redox biosensor protein Mrx1-roGFP2.

To analyze the redox states in *C. glutamicum* colonies, the 380 nm/470 nm biosensor ratios from 120 arrayed colonies of WT_Mrx1-roGFP2 and the MSH-deficient mutant strain Δ*mshC*_Mrx1-roGFP2 were determined. The Mrx1-roGFP2 biosensor consists of redox sensitive GFP2 (roGFP2) which harbors two Cys residues. Upon oxidation it forms a disulfide bond, resulting in an increase of the respective biosensor ratio (Exc. 380 nm/ Exc. 470 nm), whereas it responds in the opposite way upon reduction of the biosensor protein. As depicted in Fig. 5a, the mean values of the biosensor ratios were significantly different with 0.82 ± 0.01 and 1.52 ± 0.02 for WT_Mrx1-roGFP2 and Δ*mshC*_Mrx1-roGFP2, respectively, indicating a more oxidized state of the biosensor protein Mrx1-roGFP2 in the Δ*mshC*_Mrx1-roGFP2 strain background on agar plates (Fig. 5a). This is in accordance to previous measurements conducted in liquid cultures with biosensor ratios of 1.0 ± 0.02 (WT_Mrx1-roGFP2) and 1.52 ± 0.03 (Δ*mshC*_Mrx1-roGFP2)^16^.

Further, we validated the dynamic response and functionality of the sensor protein Mrx1-roGFP2 in colonies by applying Dithiothreitol (DTT; 100 mM) and Natriumhypochlorite (NaOCl; 100 mM) as reducing and oxidizing agents, respectively. For this, 5 μL drops were applied onto WT_Mrx1-roGFP2 and Δ*mshC*_Mrx1-roGFP2 colonies followed by real-time monitoring of the biosensor signals. Prior to the colony treatment, WT_Mrx1-roGFP2 was maintaining stable biosensor ratios between 0.76 - 0.78 followed by a strong reduction or increase of the biosensor ratio upon applying DTT or NaOCl, respectively (Fig. 5b). After five minutes of incubation, fully reduced (DTT; biosensor ratio of 0.53) and fully oxidized (NaOCl; biosensor ratio of 1.4) biosensor ratios were recorded until the end of the experiment (Fig. 5b). The treatment with PBS only temporarily induced a slight decrease of the recorded biosensor ratio but then was restored to initially recorded biosensor ratios (Fig. 5b). In contrast, Δ*mshC*_Mrx1-roGFP2 colonies revealed biosensor ratios between 1.4 - 1.45 prior to the treatment of the colonies. The addition of DTT buffer solution resulted in a strong reduction of the biosensor ratio reaching fully reduced ratios of 0.54, six to eight minutes following the treatment (Fig. 5c). As expected, treatment with NaOCl just slightly increased the ratio to 1.54 due to its almost fully oxidized state, when compared to a final biosensor ratio of 1.4 for colonies treated with PBS buffer only (Fig. 5c). To note, no alteration of the recorded fluorescence signals was observed when performing the same experiment using the *C. glutamicum* WT controls strain, indicating that the measured change of the ratiometric biosensor signals in both sensor strains can be attributed to the biosensor protein Mrx1-roGFP2 (Fig. S3).

Based on the measured biosensor ratios, the oxidation degree (OxD) of the biosensor protein Mrx1-roGFP2 was calculated according to equation 1. OxD values of 0.93 ± 0.03 and 0.33 ± 0.03 were calculated for the Mrx1-roGFP2 biosensor in Δ*mshC*_Mrx1-roGFP2 and WT_Mrx1-roGFP2 colonies on agar plates, respectively (Fig. 5d). The results are consistent with previous studies performed in shaker-flasks under non-stressed conditions where OxD values between 0.3 - 0.5 were reported for WT_Mrx1-roGFP2^16,34^. In contrast, almost fully oxidized biosensor states were reported for the MSH-deficient mutant strain (OxD = 0.8 - 0.95)^16,34^. Upon the formation of ROS, the redox-active sulfhydryl group of MSH can either scavenge free radicals directly or functions as a cofactor for antioxidant enzymes resulting in formation of oxidized mycothiol disulfide (MSSM)^31,35–38^. Accordingly, the lack of MSH elevates the intracellular ROS levels and in turn induces an auto-oxidation of the biosensor protein Mrx1-roGFP2 in Δ*mshC*_Mrx1-roGFP2. Thus, previous results and the results of this study indicate the major role of MSH for the overall redox homeostasis under aerobic growth conditions in shake-flasks cultivations but also in colonies growing on agar plates.

**Figure 5:**
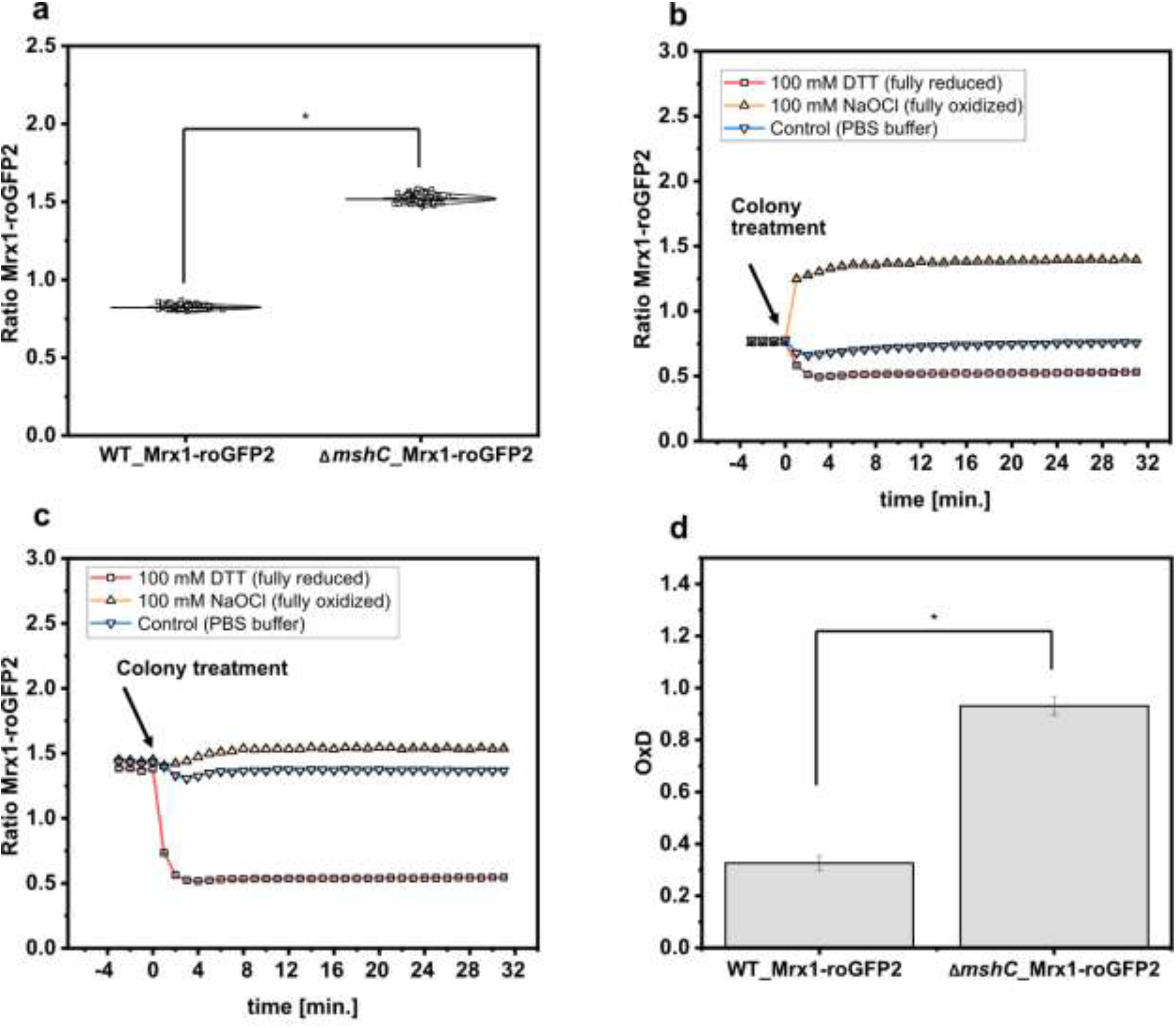
Calculated biosensor ratio (Exc. 380nm/ Exc. 470 nm) of the sensor protein Mrx1-roGFP2 in colonies of *C. glutamicum* WT_Mrx1-roGFP2 (WT_Mrx1-roGFP2) and *C. glutamicum* Δ*mshC*_Mrx1-roGFP2 (Δ*mshC*_Mrx1-roGFP2) (**a**) and real-time monitoring of the response of the ratiometric signal of the biosensor Mrx1-roGFP2 upon applying DTT (100 mM) (reducing agent), hypochlorite (100 mM) (oxidizing agent) and PBS buffer (control) onto WT_Mrx1-roGFP2 (**b**) and Δ*mshC*_Mrx1-roGFP2 colonies (**c**). Calculation of the oxidation degree (OxD) by normalizing biosensor ratios to fully oxidized and reduced states (**d**). For sensor analysis, 30 colonies from four independent agar plates and experiments were analyzed for each strain (120 colonies in total). Biosensor ratios were analyzed with one-way-ANOVA followed by Tukey’s test (^n.s^ p > 0.05; * p ≤ 0.001).

### 2.5 Ratiometric biosensor signals are independent of the colony size

Fluorescence signals derived from single fluorescent proteins need to be normalized to the OD_600_ (96-well screen) or colony size (agar screen). In contrast to single fluorescent proteins, a ratiometric fluorescent protein can be normalized to the second fluorescence signal rather than growth related parameters. This implies that a ratiometric biosensor signal should not be affected by changes of the colony size during screens on agar plates.

To test this, a dilution series (different set OD_600_) of *C. glutamicum* WT (WT), *C. glutamicum* WT_Mrx1-roGFP2 (WT_Mrx1-roGFP2) and *C. glutamicum* Δ*mshC*_Mrx1-roGFP2 (Δ*mshC*_Mrx1-roGFP2) was spotted on rectangular OmniTray plates with solidified CGXII minimal medium (1 % Glucose) (Fig. 6a). After 48 hours of incubation, the final colony size was determined for all spotted dilutions and the size reduction calculated relative to the highest applied OD_600_ (Fig. 6a). For the highest applied OD_600_ (10^1^), defined as 100 % colony size, the area was determined to be 680 ± 29, 610 ± 52 and 590 ± 49 pixels for the WT, WT_Mrx1-roGFP2 and the Δ*mshC*_Mrx1-roGFP2 colonies, respectively (Fig. 6a). Upon applying higher dilutions (stepwise 1:10), the relative colony size was reduced by approximately 10 % for each dilution step and a relative colony size of 55 ± 8 %, 61 ± 2% and 54 ± 0.4 % was reached upon applying an OD_600_ of 10^-3^ for the WT, WT_Mrx1-roGFP2 and Δ*mshC*_Mrx1-roGFP2 strain, respectively (Fig. 6a). For this dilution, the circular morphology of the colonies was impaired leading to inaccuracies with respect to the colony size determination for further dilutions (Fig. 6a). Next, the positions of the arrayed colonies from the dilution series were selected and the fluorescence intensity derived from the sensor protein Mrx1-roGFP2 analyzed. Absolute fluorescence intensities measured at 510 nm (Exc. 470 nm) revealed increased fluorescence levels in colonies of WT_Mrx1-roGFP2 (5.4 × 10^6^ – 1.3 × 10^7^ FLU) and Δ*mshC*_Mrx1-roGFP2 (4.2 × 10^6^ – 1.03 × 10^7^ FLU) strains when compared to the parental wild type strain *C. glutamicum* (1.90 – 3.90 × 10^6^ FLU) (Fig. 6b). Absolute fluorescence intensities decreased upon a reduction of the colony size (Fig. 6b). In accordance with the fluorescence analysis for mCherry or GFP in the previous sections, relative fluorescence intensities were calculated by normalizing the absolute fluorescence intensity to the perimeter of the colony. As expected, fluorescence signals normalized to the measured perimeter of the colonies resulted in stable relative fluorescence intensities until a colony size reduction of 70 % was reached (Fig. S4). Next, the second fluorescence signal (Exc. 380 nm/Em. 510 nm) was measured for all colonies and the biosensor ratio calculated (Exc. 380 nm/ Exc. 470 nm). The ratiometric biosensor signal independently of the colony size, remained stable between 1.0 – 1.1 and 1.4 – 1.5 in WT_Mrx1-roGFP2 and Δ*mshC*_Mrx1-roGFP2 colonies, respectively (Fig. 6c). As discussed in the previous section, higher biosensor ratios measured for the MSH-deficient strain are caused by the more oxidized state of the biosensor protein Mrx1-roGFP in this strain background. To note, increased ratiometric signals (between 2.5 to 4.0) for the agar surface (no colony) and for *C. glutamicum* not harboring the biosensor protein Mrx1-roGFP2 are due to low background fluorescence at 510 nm (Exc. 470 nm) but a higher background fluorescence at 510 nm when excited at 380 nm (Fig. 6c).

Taken together, the results demonstrate that the signal derived from a ratiometric biosensor is robust against variations of colony size and the morphology of colonies (Fig. 6c). This makes ratiometric FRPs a powerful tool when conducting screens of mutant libraries comprising different phenotypes (e.g. growth behavior). Furthermore, agar plate-based screens provide the advantage that the tested strains can be exposed to various conditions at the same time (replica plating) ^19,39^. Conducting screens under industrially relevant conditions (e.g. pH gradients) supported by the use of ratiometric “stress” biosensors will facilitate the development of structured metabolic models for industrially relevant organisms. This bridges metabolic engineering and bioprocess development allowing for the development of computational models suitable for both cell factory design and process optimization at industrial scales in the future ^40^.

**Figure 6:**
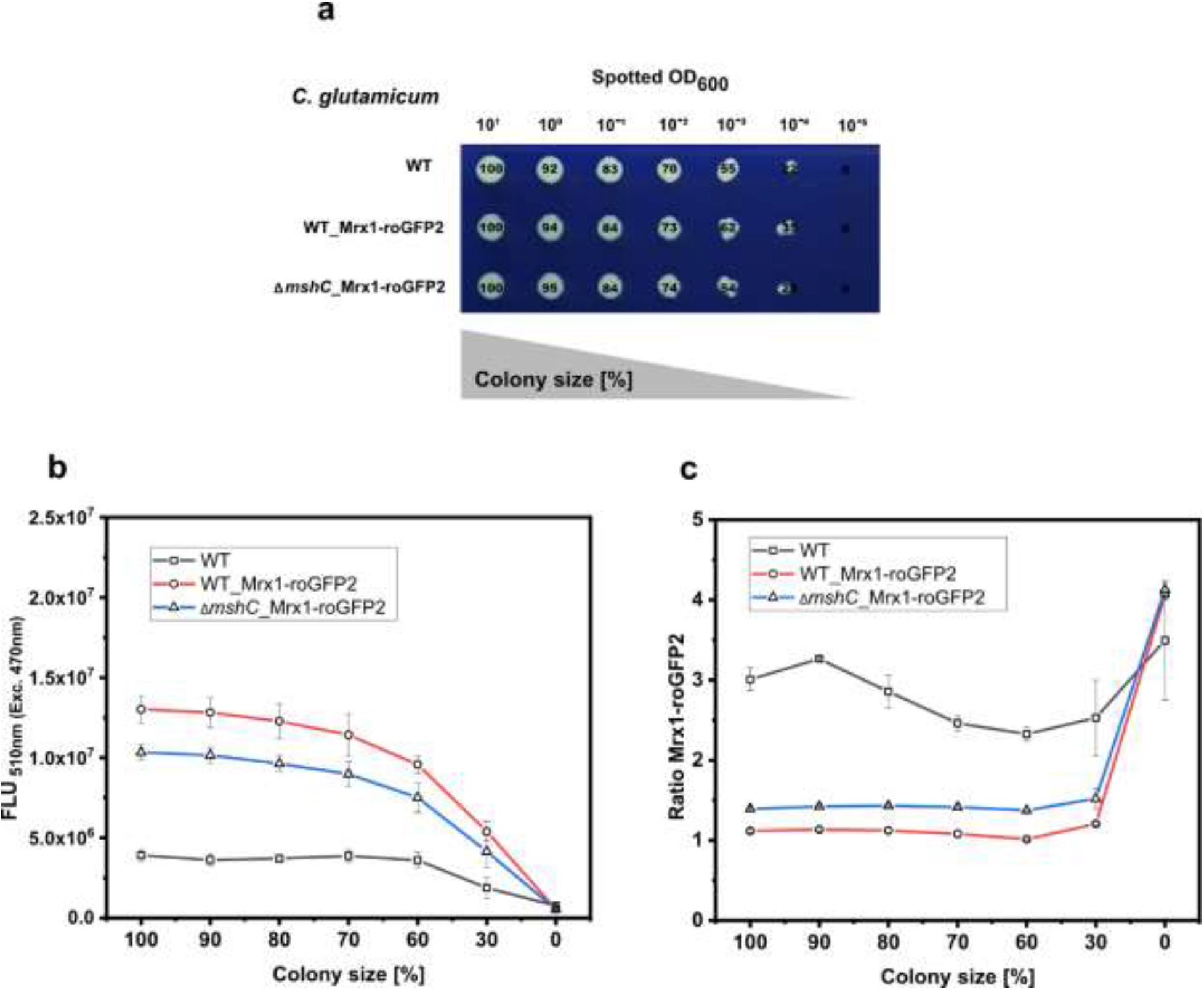
Robotic spotting of a dilution series of *C. glutamicum* WT (WT), *C. glutamicum* WT_Mrx1-roGFP2 (WT_Mrx1-roGFP2) and *C. glutamicum* Δ*mshC*_Mrx1-roGFP2 (Δ*mshC*_Mrx1-roGFP2) liquid cultures with different set optical densities measured at 600 nm (OD_600_) and determination of the colony size relative to the highest spotted OD_600_ (**a**). Absolute fluorescence intensities measured at 510 nm (Exc. 470 nm) to validate the presence of Mrx1-roGFP2 in WT_Mrx1-roGFP2 and Δ*mshC*_Mrx1-roGFP2 colonies (**b**), and the ratiometric biosensor signal of Mrx1-roGFP2 in the different colonies (**c**). Robotic spotting was performed using a replica plating Robot ROTOR. Dilution series was grown on OmniTray plates with CGXII media supplemented with 1% Glucose. Fluorescence analysis of the colonies from the dilution series was performed in a microplate reader (SpectraMax iD3). For the calculation of the ratiometric biosensor signal, the emission intensity at 510 nm was recorded upon an excitation at 380 nm and 470 nm and the former was divided by the latter (Exc. 380 nm/ Exc. 470 nm). Mean values from two independent experiments are shown.

## 3 Conclusion

Phenotypic screenings using arrayed colonies is a widely applied approach to screen strain libraries for particular variants with desired phenotypes. Genetically encoded biosensors are powerful analytical tools to support the rational decision making for strain selection as intracellular states can easily be assessed in a high-throughput manner. However, imaging analysis often limits the applicability of biosensors as the optical set-up of the utilized imaging device does not match with the properties of the fluorescent protein. The microplate reader-based system for sensor analysis established here provides the advantage of a monochromatic technology, which provides high flexibility with respect to different excitation and emission wavelengths. This novel technique has revealed high sensitivity for the detection of low fluorescence levels making it attractive for the development and optimization of genetic circuits regulating harmful targets. In addition, the monochromatic technology enables great opportunities to utilize FRPs, which application to date is often limited due to their complex ratiometric fluorescent properties. The ratiometric readout makes these types of biosensors highly robust against signal fluctuations caused by the colony size or morphology of the colony. Moreover, applications of such biosensors enables the measurement of complex intracellular states such as oxidative stress or the internal pH in real-time due to their fast dynamics. The microplate reader-based analysis established here can easily be adapted to further biosensors and combinations thereof without causing additional costs and effort to install adequate filter modules. Thus, this technique is expected to provide new possibilities for comprehensive phenotypic screenings and novel applications in metabolic engineering and systems biology.

## 4 Materials and methods

### Strains, media, and culture conditions

Bacterial strains, yeast strains, and plasmids used in this study are listed in Table 1. Cloning was carried out using *E. coli* DH5 α, cultivated in Lysogeny Broth (LB) medium^41^. *C. glutamicum* was cultivated using BHI-media (Sigma Aldrich, Germany). *E. coli* MG1655 was cultivated in SB-medium (5 g/L yeast extract; 10 g/L BactoTryptone; 100 mM NaCL; 50 mM KCl, buffered with 50 mM TRIS; pH 7.0) as recently described^8^. For preparation of agar plates, 16 g/L agar was added to the respective media. Strains carrying plasmids were cultivated in presence of kanamycin (50 μg/mL) or chloramphenicol (20 μg/mL). If required and unless stated otherwise, 1 mM IPTG was added to induce gene expression. The haploid prototrophic yeast *Saccharomyces cerevisiae* strain CEN.PK113-7D was grown in standard yeast peptone dextrose (YPD) medium containing 10 g/L yeast extract, 20 g/L peptone, and 20 g/L glucose and incubated at 30°C. For preparation of YPD agar plates 20 g/L agar was added. For selection of yeast with antibiotic resistance marker NatMX, medium was supplemented with 100mg/L nourseothricin (ClonNat, Jena Bioscience).

**Table 1:**
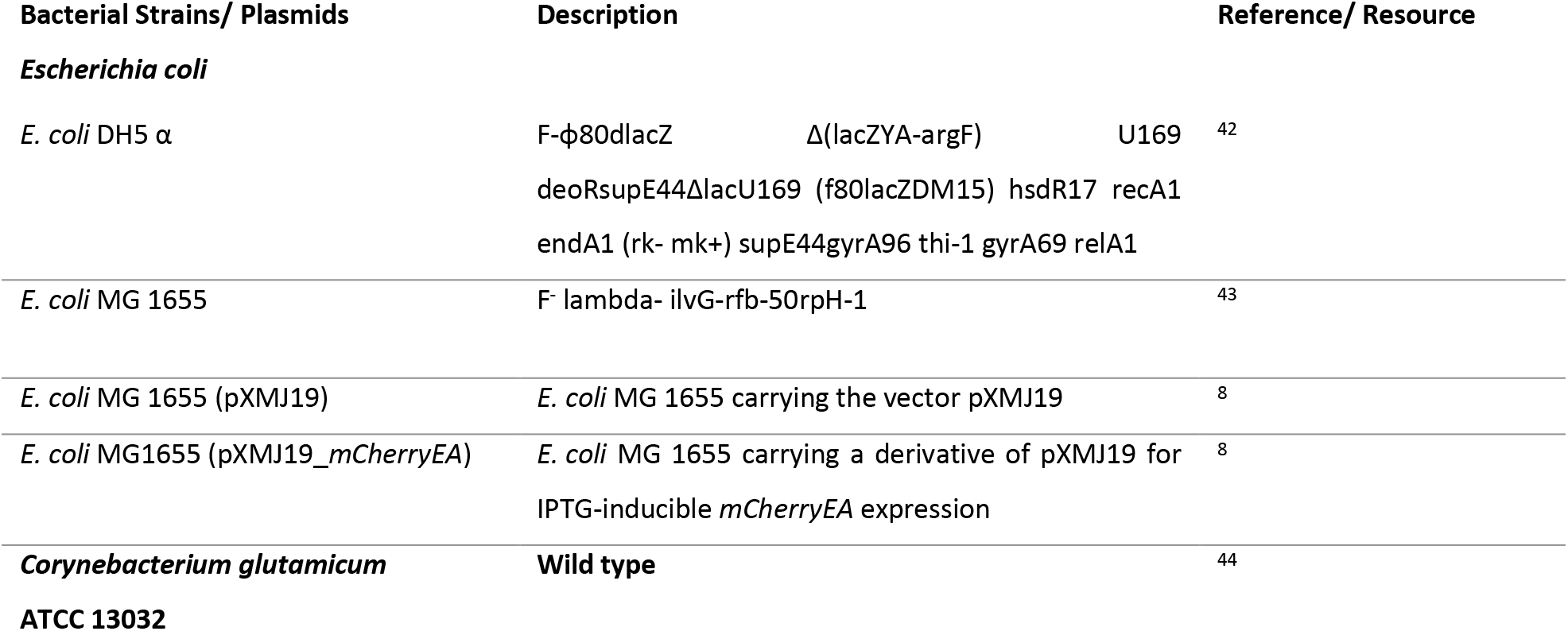
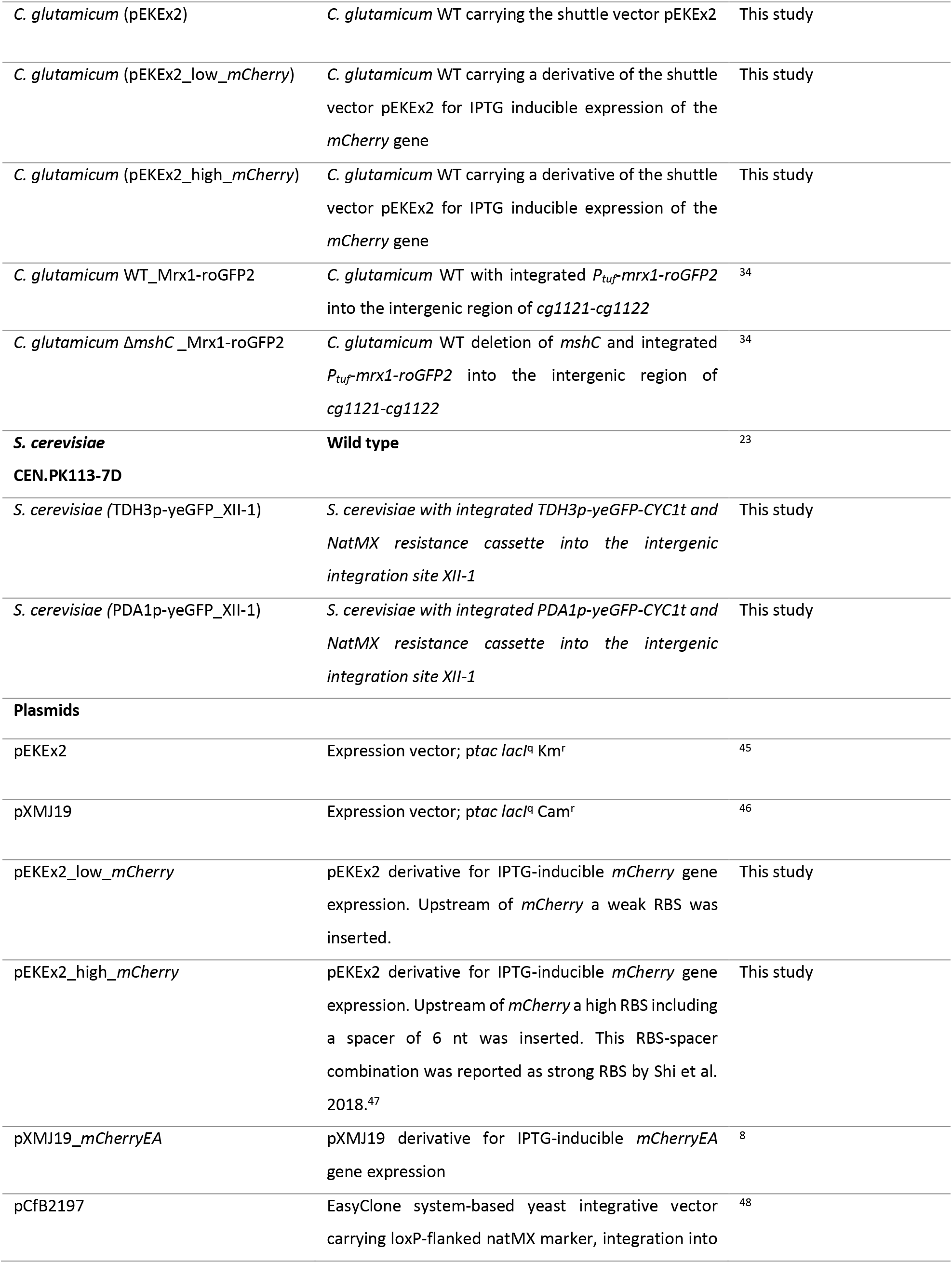
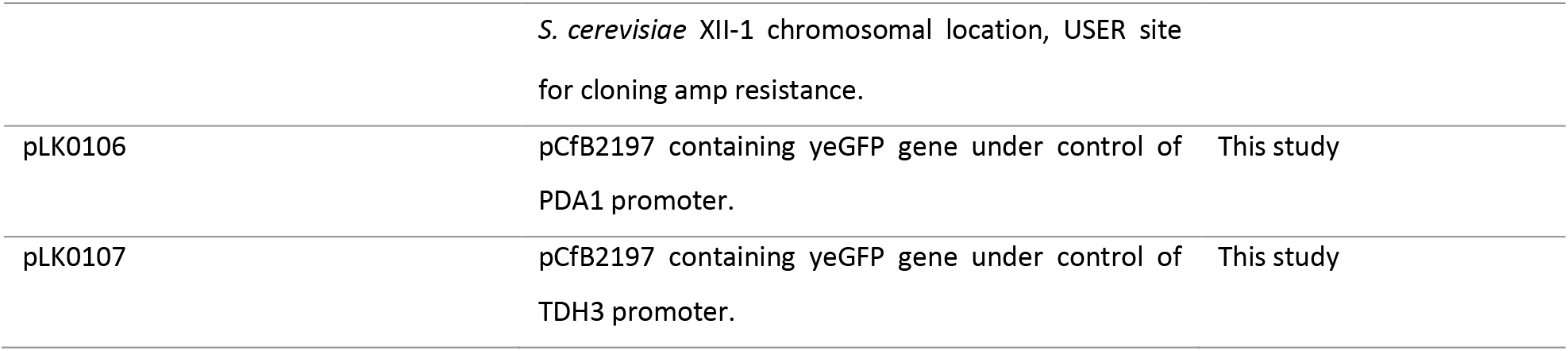
Bacterial strains, yeast strains and plasmids used in this study.

| Bacterial Strains/ Plasmids | Description | Reference/ Resource |
| --- | --- | --- |
| ***Escherichia coli*** |  |  |
| *E. coli* DH5 $\alpha$ | F-ϕ80dlacZ $\Delta$(lacZYA-argF) U169<br>deoRsupE44ΔlacU169 (f80lacZDM15) hsdR17 recA1<br>endA1 (rk- mk+) supE44gyrA96 thi-1 gyrA69 relA1 | 42 |
| *E. coli* MG 1655 | F- lambda- ilvG-rfb-50rpH-1 | 43 |
| *E. coli* MG 1655 (pXMJ19) | *E. coli* MG 1655 carrying the vector pXMJ19 | 8 |
| *E. coli* MG1655 (pXMJ19_ *mCherryEA*) | *E. coli* MG 1655 carrying a derivative of pXMJ19 for<br>IPTG-inducible *mCherryEA* expression | 8 |
| ***Corynebacterium glutamicum***<br>**ATCC 13032** | **Wild type** | 44 |
| *C. glutamicum* (pEKEx2) | *C. glutamicum* WT carrying the shuttle vector pEKEx2 | This study |
| *C. glutamicum* (pEKEx2_low_mCherry) | *C. glutamicum* WT carrying a derivative of the shuttle vector pEKEx2 for IPTG inducible expression of the *mCherry* gene | This study |
| *C. glutamicum* (pEKEx2_high_mCherry) | *C. glutamicum* WT carrying a derivative of the shuttle vector pEKEx2 for IPTG inducible expression of the *mCherry* gene | This study |
| *C. glutamicum* WT_Mrx1-roGFP2 | *C. glutamicum* WT with integrated *P<sub>tuf</sub>-mrx1-roGFP2* into the intergenic region of *cg1121-cg1122* | <sup>34</sup> |
| *C. glutamicum* Δ*mshC*_Mrx1-roGFP2 | *C. glutamicum* WT deletion of *mshC* and integrated *P<sub>tuf</sub>-mrx1-roGFP2* into the intergenic region of *cg1121-cg1122* | <sup>34</sup> |
| ***S. cerevisiae***<br>**CEN.PK113-7D** | **Wild type** | <sup>23</sup> |
| *S. cerevisiae* (TDH3p-yeGFP_XII-1) | *S. cerevisiae* with integrated *TDH3p-yeGFP-CYC1t* and *NatMX* resistance cassette into the intergenic integration site *XII-1* | This study |
| *S. cerevisiae* (PDA1p-yeGFP_XII-1) | *S. cerevisiae* with integrated *PDA1p-yeGFP-CYC1t* and *NatMX* resistance cassette into the intergenic integration site *XII-1* | This study |
| **Plasmids** |  |  |
| pEKEx2 | Expression vector; *ptac lacI<sup>q</sup>* Km<sup>r</sup> | <sup>45</sup> |
| pXMJ19 | Expression vector; *ptac lacI<sup>q</sup>* Cam<sup>r</sup> | <sup>46</sup> |
| pEKEx2_low_mCherry | pEKEx2 derivative for IPTG-inducible *mCherry* gene expression. Upstream of *mCherry* a weak RBS was inserted. | This study |
| pEKEx2_high_mCherry | pEKEx2 derivative for IPTG-inducible *mCherry* gene expression. Upstream of *mCherry* a high RBS including a spacer of 6 nt was inserted. This RBS-spacer combination was reported as strong RBS by Shi et al. 2018.<sup>47</sup> | This study |
| pXMJ19_mCherryEA | pXMJ19 derivative for IPTG-inducible *mCherryEA* gene expression | <sup>8</sup> |
| pCfB2197 | EasyClone system-based yeast integrative vector carrying loxP-flanked natMX marker, integration into | <sup>48</sup> |
|  | *S. cerevisiae* XII-1 chromosomal location, USER site<br>for cloning amp resistance. |  |
| pLK0106 | pCfB2197 containing yeGFP gene under control of PDA1 promoter. | This study |
| pLK0107 | pCfB2197 containing yeGFP gene under control of TDH3 promoter. | This study |

### Strain construction

The two plasmids pEKEx2_low_*mCherry* and pEKEx2_high_*mCherry* were assembled *via* Gibson cloning using the Gibson Assembly^®^ Master Mix (NEB, USA) according to the manufacturer’s instructions. For his purpose pEKEx2 was first linearized using SacI (NEB, USA). The gene for mCherry was amplified by PCR using the primer pairs low_*mCherry_fw* and low_mCherry_rev (Table S1) or high_*mCherry*_fw and high_*mCherry*_rev for pEKEx2_low_*mCherry* and pEKEx2_high_*mCherry*, respectively. All plasmids were introduced into competent *E. coli* DH5α and analyzed via sequencing prior to further use. Transformation of electrocompetent *C. glutamicum* cells and strain validation were performed as described previously^49^.

To construct the two plasmids pLK0106 and pLK0107 for yeast, DNA fragments constituting the weak PDA1 promoter or the strong TDH3 promoter ^24^ and the yeGFP open reading frame ^50^ were synthesized as double-stranded gene fragments (Twist Bioscience). The integrative vector pCfB2197 from the EasyClone 2.0 toolkit was used for plasmid construction, which allowed for selection in prototrophic strains ^48^. Briefly, the vector was linearized by digestion with *AsiSI* (Life Technologies) restriction endonuclease and nicked with *Nb.BsmI* (New England BioLabs). Synthetic DNA fragments were PCR-amplified - using forward primer FW_USER_TDH3 or FW_USER_PDA1 combined with reverse primer RV_USER_yeGFP - and subsequently cloned into the linearized vector backbone by uracil-excision based (USER) cloning technique ^51,52^. Plasmids were transformed into *E. coli* for storage and amplification. Correct plasmid assembly was verified by PCR and Sanger sequencing (Eurofins Genomics) using primers ADH1_test_fw and CYC1_test_rv. Constructed integrative vectors (pLK0106 and pLK0107) were *NotI* (Life Technologies) digested to capture the linear fragments for integration. *S. cerevisiae* was transformed as previously described ^53^. For genetic integration of cassettes, 1 μg of linear DNA was transformed into yeast for integration into the XII-1 chromosomal integration site ^54^. Cells were plated onto selective YPD plates and colonies were re-streaked on selective plates. Genomic DNA was extracted as previously described ^55^, and integration verified by PCR using XII-1_up and XII-1_down. All oligonucleotides used in this study can be found in the supplementary material (Table S1).

### Fluorescence analysis

Fluorescence measurements of liquid cultures were conducted in black clear-bottomed 96-well microplates (Greiner Bio-One, Austria) using a SpectraMax iD3 multi-mode plate reader (Molecular Devices LLC, U.S.A). For fluorescence analysis of the fluorescent protein mCherry, endpoint measurements were recorded by setting the excitation wavelength at 580 nm and the emission wavelength at 620 nm. For liquid cultures, fluorescence intensities were normalized to the optical density measured at 600 nm (OD_600_) using transparent flat-bottomed 96-well microplates (Greiner bio-one B.V., Netherlands). Prior to measurements of liquid cultures, cells were washed and four times concentrated.

For fluorescence analysis of the pH-sensitive protein mCherryEA, excitation scans were recorded by setting the excitation wavelength between 400 nm and 590 nm and the emission wavelength at 630 nm. For ratiometric analysis of the biosensor signal, the emission maxima obtained upon an excitation at 454 nm and 580 nm were taken and the corresponding biosensor ratio was calculated by dividing the former emission intensity by the latter as recently described^8^. For fluorescence analysis of the redox biosensor protein Mrx1-roGFP2, the calculation of Exc. 380 nm/ Exc. 470 nm (Em. 510 nm) was used for the determination of the biosensor ratio as recently described^16^. For determination of the biosensor oxidation degree (OxD), biosensor ratios from untreated samples were normalized to fully reduced (100 mM Dithiothreitol (DTT) in PBS buffer; pH 7.0) or oxidized (100 mM Natriumhypochlorite (NaOCl) in PBS buffer, pH 7.0) samples according to equation 1. Here, I380_sample_ and I470_sample_ represent the measured fluorescence intensities received for an excitation at 380 nm and 470 nm, respectively. Fully reduced and oxidized controls are given by I380_red_, I470_red_ and I380_ox_, I470_ox_, respectively.

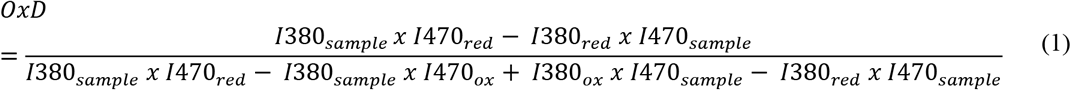

### Microplate reader-based analysis of sensor signals in colonies on agar plates

Prior to robotic spotting of cell cultures, wells of 96-well microtiter plates (Greiner bio-one B.V., Netherlands) were filled with 200 μL overnight culture adjusted to an OD_600_ of 1 in the respective media. For dilution series, the OD_600_ was set to 10 and 1:10 dilutions prepared in microtiter plates. The working plate (96-well plate) was used as a source plate for robotic spotting using a ROTOR HDA benchtop robot (Singer Instruments, United Kingdom) on rectangular OmniTray plates (Singer Instruments, United Kingdom) with solidified medium on the target plate as recently described^8^. The OmniTray plates were prepared by using 50 mL of the respective media supplemented with appropriate antibiotic and IPTG for biosensor expression if required. Agar plates with spotted cell cultures in an arrayed format were cultivated for 24 hours (*E. coli* and *S. cerevisiae* strains) or 48 hours for *C. glutamicum* strains if not otherwise stated. Fluorescence analysis was performed using a SpectraMax iD3 multi-mode plate reader (Molecular Devices LLC, U.S.A). Prior to fluorescence analysis, the lid from the OmniTray plate was removed. A 96-well plate scheme was selected for the measurement mode and the wells selected for targeting the respective colonies arrayed in a 96-well scheme on the agar plates. Measurement height was set to 5 mm and measurements were performed at ambient temperature.

### Fluorescence imaging of arrayed colonies

Fluorescence imaging of microbial colonies on agar plates was carried out using the photo-documentation system FUSION FX (Vilber Lourmat, France) and the FUSION FX EVOLUTION-CAPT software (Vilber Lourmat) for image analyses. For fluorescence analysis of the fluorescent protein mCherry, the FUSION FX was equipped with a capsule for excitation at 530 nm and an emission filter at 595 nm. For yeGFP analysis a capsule for excitation at 480 nm and a filter for emission 530 nm were used. Exposure time was set to 800 msec if not stated otherwise. Biosensor signals from the pH-sensitive protein mCherryEA were measured using the FUSION FX as recently described ^8^. For fluorescence analysis using the Phenobooth (Singer Instruments, United Kingdom), the blue channel was selected for excitation (470 nm) and the emission intensity measured using a GFP filter (527 nm/ 20 nm).

### White light imaging and determination of colony sizes

White light images were captured using the Phenobooth (Singer Instruments, United Kingdom). If required, images were processed prior to analyzing the images using Cellprofiler 4 (version 4.0.7) ^56^. In order to determine colony sizes, the respective images were analyzed *via* the Python toolbox Pyphe ^57^.

### Statistical analysis

Biosensor signals were analyzed using one-way variance (ANOVA) followed by Tukey’s test. The respective analysis was performed using Python 3 ^58^, and Pandas ^59^ was used to handle data frames. ANOVA was performed using the ols() and anova_lm() function of the statsmodels package ^60^. Tukey’s test was performed using tukey_hsd() function of the bioinfokit package ^61^. Differences were considered significant when p-value < 0.05. All data were plotted and visualized using the software Origin.

## Abbreviations

CTAB: Cetyltrimethylammonium bromide
DTT: Dithiothreitol
EV: Empty vector
FRP: Fluorescent reporter protein
GFP: Green fluorescent protein
IPTG: Isopropyl β-d-1-thiogalactopyranoside
OD: Optical density
OxD: Oxidation degree
RBS: Ribosomal binding site
RFLU: Relative fluorescence units
roGFP2: Redox sensitive GFP2
TFB: Transcription factor-based biosensor
WT: Wild type

## Credit author statement

**Fabian S. F. Hartmann:** Conceptualization; Data curation, Formal analysis, Investigation, Methodology, Validation, Visualization, Writing of original draft; Writing-review & editing; **Tamara Weiß:** Conceptualization; Data curation, Formal analysis, Investigation, Methodology, Validation, Visualization, Writing of original draft; Writing-review & editing; **Louise L. B. Kastberg:** Formal analysis, Investigation, Writing-review & editing; **Christopher T. Workman:** Funding acquisition, Writing-review & editing; **Gerd M. Seibold:** Conceptualization; Funding acquisition; Supervision; Writing-original draft; Writing-review & editing.

## Availability of data and materials

All data generated and analyzed during this study are included in this article and its additional files. Raw datasets are available from the corresponding author on reasonable request.

## Acknowledgment

We would like to thank the Fermentation Core at DTU Bioengineering for excellent technical support.

