## SupplementaryData for "Precise and versatile microplate reader-based analyses of biosensor signals from arrayed microbial colonies"

Fabian Stefan Franz Hartmann[1†], Tamara Weiß[1†], Louise La Barbera Kastberg[1], Christopher T. Workman[1], Gerd Michael Seibold[1*]

[1] Department of Biotechnology and Biomedicine, Section for Synthetic Biology, Technical University of Denmark, Kongens Lyngby, Denmark

The supplementary material comprises one table and four figures:

**Table S1:** All oligonucleotides used in this study.

**Figure S1:** Fluorescence analysis of the fluorescent protein mCherry in different *C. glutamicum* colonies using the Vilber FUSION FX Gel Doc imaging system with different set exposure times.

**Figure S2:** Absorbance well- scan of colonies arrayed in a 96- well scheme.

**Figure S3:** Dynamic response of *C glutamicum* WT upon applying different agents on colonies.

**Figure S4:** Relative fluorescence intensities determined for different colony sizes of *C. glutamicum* WT, *C. glutamicum* WT_Mrx1-roGFP2 and *C. glutamicum* Δ*mshC*_Mrx1-roGFP2.

*Table S1: All oligonucleotides used in this study.*

| Oligonucleotide | Sequence (5’-3’)[1] | Reference | Description |
|---|---|---|---|
| low_*mCherry*_fw | ggatccccgggtaccgagct*AAAGGAGGCCCTTCAG*ATG | This study | Forward primer with low RBS and amplification of *mCherry* gene |
| low_*mCherry*_rev | acggccagtgaattcgagctTTACTTGTACAGCTCGTCC | This study | Reverse primer with low RBS and amplification of *mCherry* gene |
| high_*mCherry*_fw | ggatccccgggtaccgagct*gaaaggagaggattg*ATGGTGAGCAAGGGCGAG | This study | Forward primer with high RBS and amplification of *mCherry* gene |
| high_*mCherry*_rev | acggccagtgaattcgagctgagctcTTACTTGTACAGCTCGTCCATG | This study | Reverse primer with high RBS and amplification of *mCherry* gene |
| XII-1_up | CTGGCAAGAGAACCACCAAT | (Stovicek et al., 2015) | Forward primer for integration verification in S. cerevisiae site XII-1. |
| XII-1_down | GGACGACAACTACGGAGGAT | (Stovicek et al., 2015) | Reverse primer for integration verification in S. cerevisiae site XII-1. |
| ADH1_test_fw | GAAATTCGCTTATTTAGAAGTGTC | (Stovicek et al., 2015) | verification of a gene expression cassette cloning into EasyClone vectors |
| CYC1_test_rv | CTCCTTCCTTTTCGGTTAGAG | (Stovicek et al., 2015) | verification of a gene expression cassette cloning into EasyClone vectors |
| FW_USER_TDH3 | CGTGCGAUTCATTATCAATACTCGCCATTT | This study | Reverse primer for amplification of constructs with yeGFP to generate USER overhangs for AsiSI + Nb.BsmI USER cloning |

| | | | |
|---|---|---|---|
| FW_USER_PDA1 | CGTGCGAUTGAAATTCAAAACTCTCCAGAC | Thus study | Forward primer for amplification of constructs with S. cerevisiae TDH3 promoter to generate USER overhangs for AsiSI + Nb.BsmI USER cloning |
| RV_USER_yeGFP | CACGCGAUTTATTTGTACAATTCATCCATACC | This study | Forward primer for amplification of constructs with S. cerevisiae PDA1 promoter to generate USER overhangs for AsiSI + Nb.BsmI USER cloning |

[1]Capital letters: annealing sites, italic letters: RBS, underlined letters: spacer

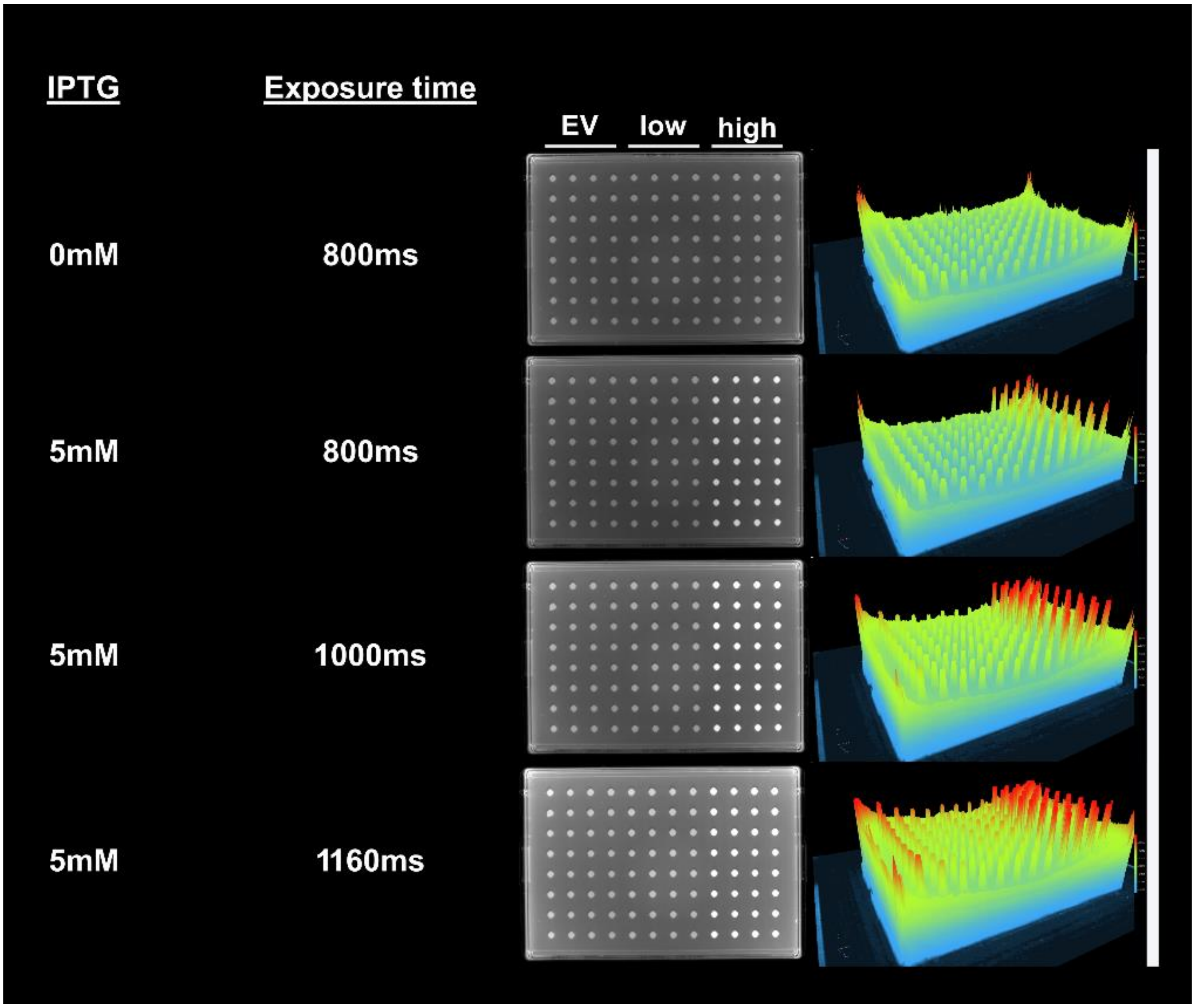


Figure S1: Fluorescence analysis of the fluorescent protein mCherry in *C. glutamicum* (pEKEx2) (EV), *C. glutamicum* (pEKEx2_low_*mCherry*) (low) and *C. glutamicum* (pEKEx2_high_*mCherry*) (high) colonies on BHI agar plates (0 mM and 5 mM IPTG) via imaging using the FUSION FX Gel Doc imaging system (Vilber). For fluorescence analysis of the fluorescent protein mCherry, the FUSION FX was equipped with a capsule for excitation (530 nm) and an emission filter (595 nm). Different exposure times were tested. Upon increasing the exposure time (1000ms or 1160ms), the signals obtained from the colonies equipped with the pEKEx2_high_mCherry construct reached saturation.

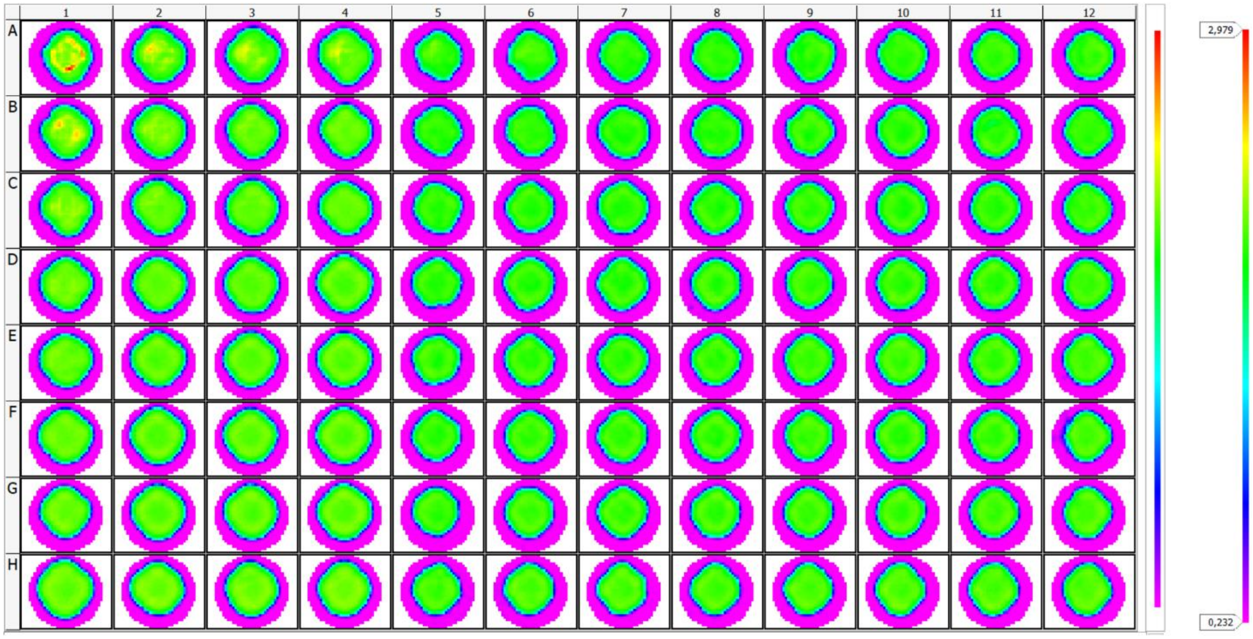


Figure S2: Absorbance well- scan (30 x 30) of an OmniTray agar plate with arrayed colonies of *C. glutamicum* with different sizes. Measured absorbance is visualized with pink, yellow and red colored pixels indicating increasing absorbance intensities, respectively. With this scanning mode, the surface of the agar (pink) can be distinguished from colonies and by this the location of the colony determined. Irrespectively of the colonies´ position, all colonies were found to be centered for the scanned area, corresponding to the well position of a 96- well microtiter plate. Absorbance scan was conducted using a microplate reader (CLARIOstar$^{Plus}$; BMG LABTECH, Germany).

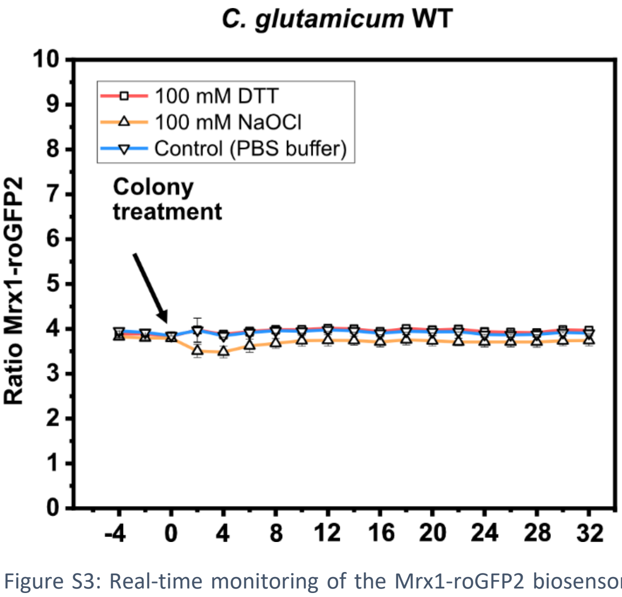


Figure S3: Real-time monitoring of the Mrx1-roGFP2 biosensor ratio ((Exc. 380 nm/ Exc. 470 nm) and Em. 510 nm) in arrayed colonies of *C. glutamicum* WT before and after applying 5 µL PBS, DTT and Hypochlorite (NaOCl) as control, reducing agent and oxidizing agent, respectively. Fluorescence measurements were conducted in a microplate reader (SpectraMax). Colonies were arrayed on OmniTray plates using BHI- medium. Mean values are shown from three independent experiments.

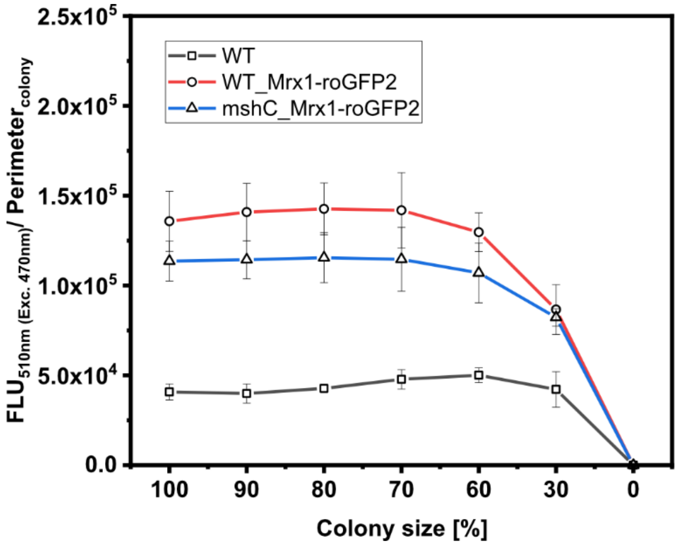


Figure S4: Robotic spotting of a dilution series of *C. glutamicum* WT (WT), *C. glutamicum* WT_Mrx1-roGFP2 (WT_Mrx1-roGFP2) and *C. glutamicum* Δ*mshC*_Mrx1-roGFP2 (Δ*mshC*_Mrx1-roGFP2) liquid cultures with different set optical densities measured at 600 nm ($OD_{600}$). Absolute fluorescence intensities measured at 510 nm (Exc. 470 nm) were normalized to the respective size (perimeter) of the colonies. Robotic spotting was performed using a replica plating Robot ROTOR. Dilution series was grown on OmniTray plates with CGXII media supplemented with 1% Glucose. Fluorescence analysis of the colonies from the dilution series was performed in a microplate reader (SpectraMax). Mean values from two independent experiments are shown. Mean values are shown from two independent experiments.
